# Disruption of the nuclear localization signal in RBM20 is causative in dilated cardiomyopathy

**DOI:** 10.1101/2022.12.08.519616

**Authors:** Yanghai Zhang, Zachery R. Gregorich, Yujuan Wang, Camila Urbano Braz, Jibin Zhang, Yang Liu, Peiheng Liu, Nanyumuzi Aori, Timothy A. Hacker, Henk Granzier, Wei Guo

**Author notes:** To whom correspondence should be addressed: Dr. Wei Guo, 1933 Observatory Dr., Madison, WI 53706, United States. Equal contribution (Yanghai Zhang, Zachery R. Gregorich).

## Abstract

Human patients carrying genetic mutations in RNA binding motif 20 (RBM20) develop a clinically aggressive dilated cardiomyopathy (DCM). RBM20 is a splicing factor with two canonical domains, an RNA recognition motif (RRM) and an arginine-serine rich (RS) domain. RRM loss-of-function disrupts the splicing of RBM20 target transcripts and leads to systolic dysfunction without overt DCM, while mutations in the RS domain precipitate DCM. We show that mice lacking the RS domain (*Rbm20*^*ΔRS*^) manifest DCM with mis-splicing of RBM20 target transcripts. We found that RBM20 is mis-localized in *Rbm20*^*ΔRS*^ mice but not in mice lacking the RRM, which are also deficient in RBM20 splicing. We determine that the RS domain, not other domains including the RRM, is critical for RBM20 nuclear import and define the core nuclear localization signal (NLS) within this domain. Mutation analysis of phosphorylation sites within the RS domain indicate that phosphorylation is dispensable for RBM20 nuclear import. Collectively, our findings establish disruption of the NLS in RBM20 as a causative mechanism in DCM through nucleocytoplasmic transport.

## INTRODUCTION

Dilated cardiomyopathy (DCM) is a common cause of heart failure and has an estimated prevalence of 1 in 250 individuals in the general population.^1^ Mutations in the *TTN* gene, which encodes the giant sarcomeric protein titin, account for approximately 25% of familial cases of idiopathic DCM and 18% of sporadic cases, making *TTN* mutations the most common cause of DCM.^2^ Yet, the genetics of DCM are complex with mutations in over 60 genes already linked to this devastating disease.^3^ Among the myriad DCM-linked genes that have been identified, RNA binding motif 20 (RBM20) is unique in being one of only two known splicing factors that, when mutated, leads to the development of DCM in humans and animal models.^4-8^ Autosomal dominant mutations in RBM20 account for approximately 3% of familial DCM cases.^9,10^ RBM20 is a muscle-specific splicing factor that belongs to the serine arginine (SR) protein family.^11^ Like other SR family proteins, RBM20 contains a C-terminal domain rich in arginine and serine dipeptides, termed the RS domain, as well as an N-terminal RNA recognition motif (RRM). In general, the RRM(s) of SR proteins confers RNA-binding specificity while the RS domain mediates protein-protein interactions between SR proteins and other components of the splicing machinery, as well as nuclear localization.^12-14^

RBM20 knockout rats and mice developed a DCM-like phenotype characterized by dilation of the left ventricle (LV), fibrosis, and an increased incidence of arrhythmia.^15,16^ It was originally accepted that this phenotype arises as a consequence of the mis-splicing of RBM20 target genes.^15,16^ Yet, this splicing-centric view was challenged when mice expressing RBM20 lacking the RRM domain (*Rbm20*^*ΔRRM*^) were found to exhibit disrupted splicing of RBM20 target genes and systolic dysfunction without overt DCM phenotype.^17^ Similarly, it was recently demonstrated that mice harboring the I536T variant, which is located within the RRM, develop neither DCM nor cardiac dysfunction despite altered splicing of RBM20 target transcripts.^18^ Given that the RBM20 RS domain is a hotspot for DCM-linked mutations (along with the RRM and E-rich regions),^19,20^ in this study we sought to (1) determine the function of the RBM20 RS domain in DCM and (2) examine the underlying mechanism(s) of its regulation in DCM-related mis-localization.

## RESULTS

### Generation and characterization of *Rbm20*^*ΔRS*^ mice

To determine the function of the RS domain in RBM20, we generated mice expressing RBM20 lacking this domain. In humans, the RBM20 RS domain extends from amino acid residue 632 to 666,^15^ which corresponds to amino acids 634-657 in the mouse (Figure 1A). Using CRISPR/Cas9 genome editing, the sequence corresponding to amino acid residues 633-666 in mouse RBM20 was deleted in-frame to generate *Rbm20*^*ΔRS*^ mice (Figure 1B). Deletion of the target sequence (102 bp) in *Rbm20*^*ΔRS*^ mice was confirmed by PCR and deep sequencing using an Illumina MiSeq System (Figure 1B and Figure S1). *Rbm20*^*ΔRS*^ mice were viable, fertile, and born at expected Mendelian and sex ratios. In contrast to previously described mouse lines carrying pathogenic RBM20 mutations,^6,8^ male and female *Rbm20*^*ΔRS*^ mice did not exhibit early mortality (Figure 1C); although we did observe that *Rbm20*^*ΔRS*^ dams tended to die at a younger age than their un-mated counterparts (data not shown). In general, hearts from 4-month-old *Rbm20*^*ΔRS*^ mice of both sexes were similar in size to those of age- and sex-matched WT controls, but lacked the structural integrity of WT hearts, appearing flimsy and deflated by comparison (Figure 1D). Body weight (BW), heart weight (HW), and HW/BW ratios did not differ significantly between male and female *Rbm20*^*ΔRS*^ mice and age-matched WT controls of the same sex (Figure 1E-G). Histological examination revealed dilation of the LV in both male and female *Rbm20*^*ΔRS*^ mice by 2-months-of-age (Figure 1H). There was also a trend towards increased fibrosis in the hearts of 4-month-old female, but not male, *Rbm20*^*ΔRS*^ mice relative to that in WT mice of the same age and sex (Figure S2).

**Figure 1.**
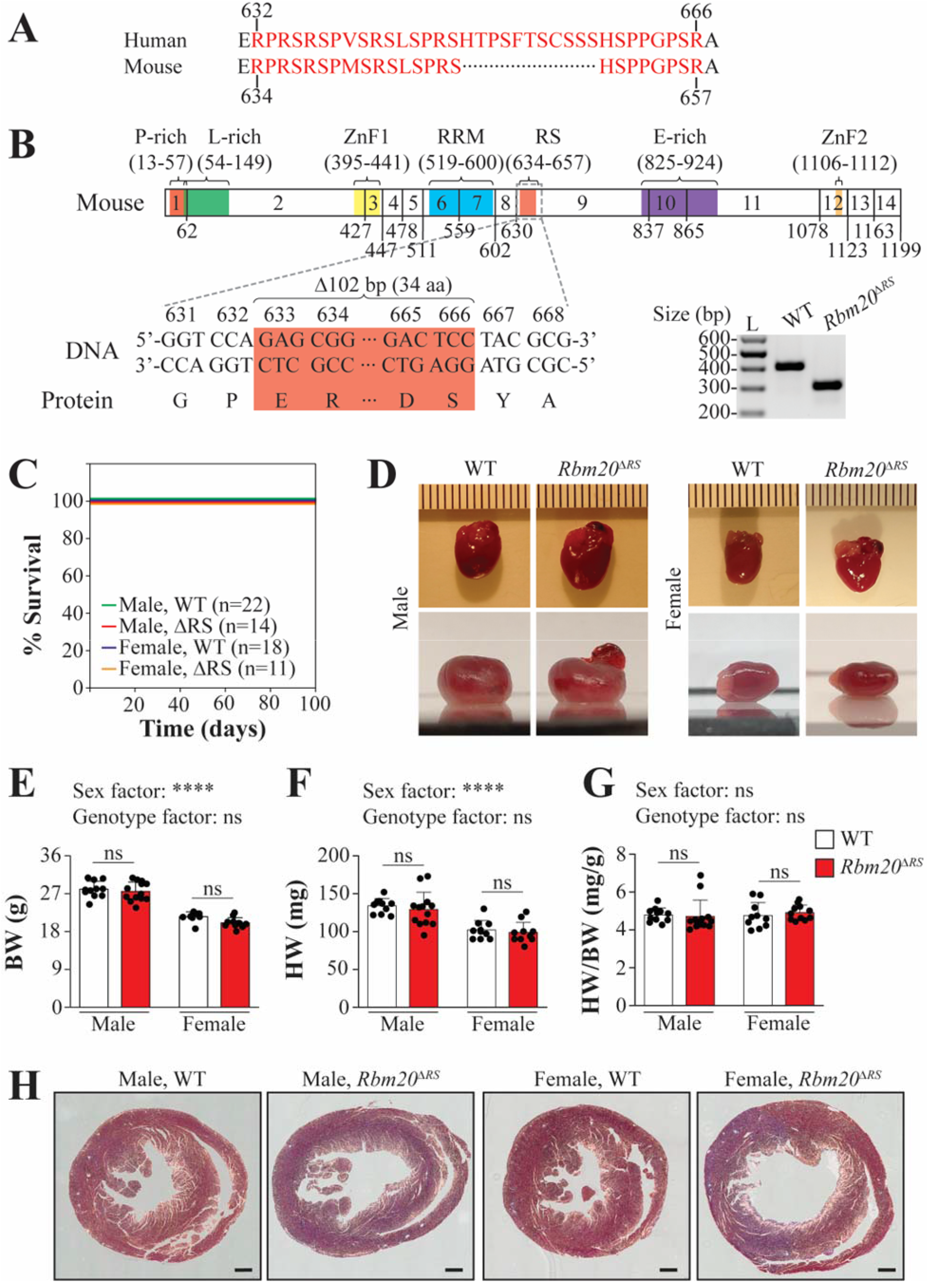
Characterization of *Rbm20*^*ΔRS*^ mice. **A**, Sequence alignment showing the sequence of the RS domain in human RBM20 and the corresponding sequence in mouse (amino acids 634-657). **B**, Schematic showing the domain structure of mouse RBM20 and genetic manipulation to produce *Rbm20*^*ΔRS*^ mice. A 102 bp stretch was removed using CRISPR/Cas9 genome editing resulting in RBM20 lacking amino acid residues 633-666. **C**, Kaplan-Meier survival curves for male and female WT and *Rbm20*^*ΔRS*^ mice. **D**, Gross morphological characterization of hearts from 4-month-old male and female WT and *Rbm20*^*ΔRS*^ mice. Ruler spacing is 0.1 cm between ticks. **E, E-F**, Bar graphs showing BW (**E**), HW (**F**), and HW/BW ratios (**G**) for 4-month-old male and female WT and *Rbm20*^*ΔRS*^ mice. Data are shown as mean ± standard deviation. Dots represent measurements from individual animals. Two-way ANOVA with the Šídák method for multiple comparisons was performed to analyze the effect of sex and genotype on each of the aforementioned parameters. The interaction between sex and genotype was not significant for any parameter. ns, not significant; \*\*\*\**p*<0.0001. **H**, Representative Masson’s trichrome-stained heart sections from 2-month-old male and female WT and *Rbm20*^*ΔRS*^ mice. Scale bars are 500 μm.

### *Rbm20*^*ΔRS*^ mice develop a DCM-like phenotype characterized by chamber dilation and systolic dysfunction

To evaluate the effects of RBM20 RS domain deletion on cardiac function, male and female *Rbm20*^*ΔRS*^ mice were evaluated by non-invasive echocardiography at 4-months-of-age. Consistent with histological examination, echocardiography confirmed that the inner diameter of the LV was significantly increased at the end of systole (LVID;s) and diastole (LVID;d) in *Rbm20*^*ΔRS*^ mice of both sexes compared to age-matched WT controls of the same sex (Figure 2A-2B and Table S3). In line with this, the end systolic (ESV) and diastolic (EDV) volumes were significantly increased in the hearts of *Rbm20*^*ΔRS*^ mice of both sexes (Figure 2C-2D and Table S3). Systolic function was impaired in both male and female *Rbm20*^*ΔRS*^ mice, which had significantly reduced stroke volumes (SVs), ejection fractions (EFs), and fractional shortening (FS) compared to age- and sex-matched WT controls (Figure 2E-2G and Table S3). Cardiac output (CO) was significantly decreased in female, but not male, *Rbm20*^*ΔRS*^ mice in comparison to same-sex WT controls (Figure 2H and Table S3). Taken together, these results demonstrate that *Rbm20*^*ΔRS*^ mice develop a DCM-like phenotype that is sexually dimorphic with females developing a slightly more severe phenotype than their male counterparts (Figure 2 and Table S3).

**Figure 2.**
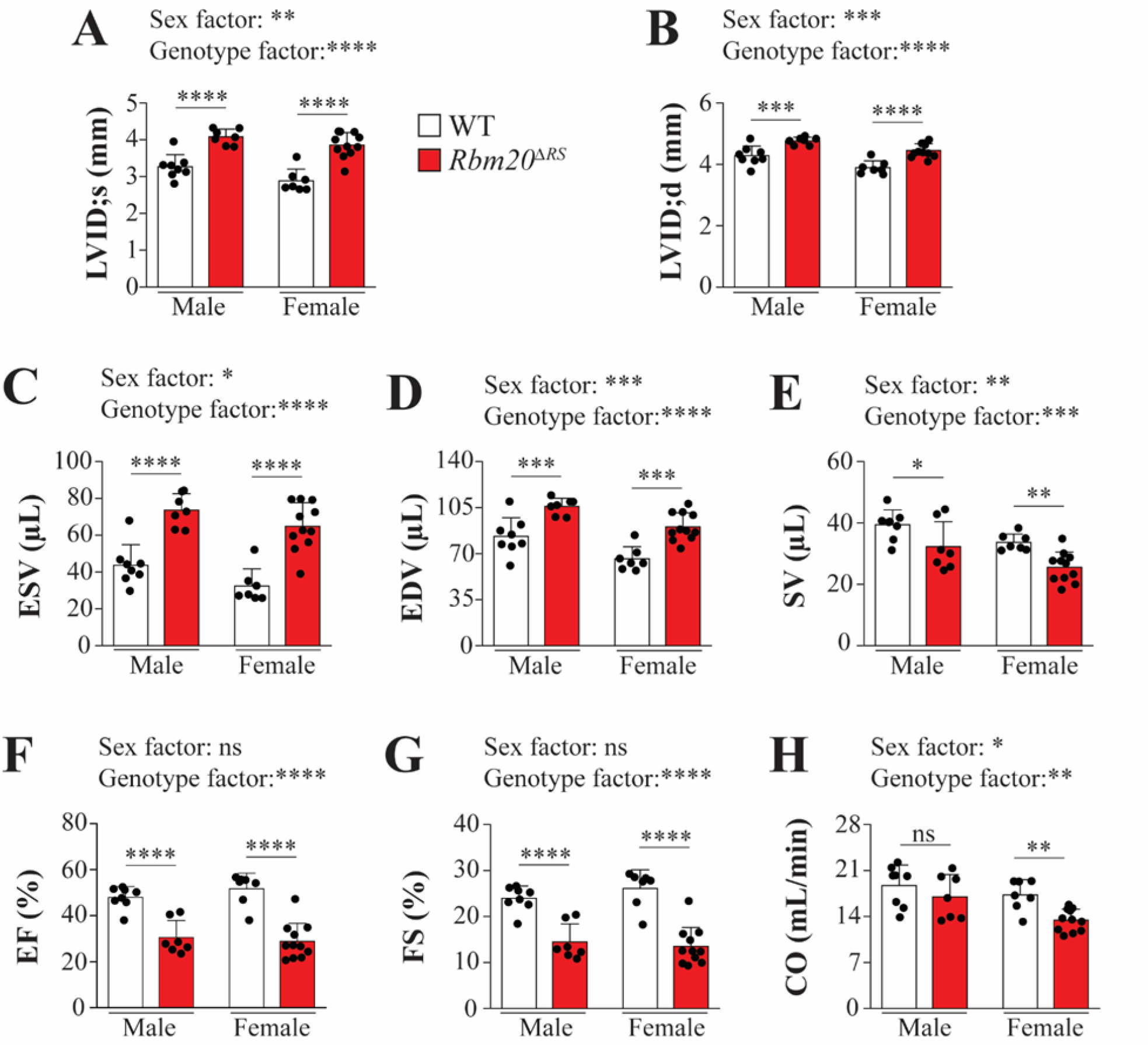
Male and female *Rbm20*^*ΔRS*^ mice develop a DCM-like phenotype. **A-H**, Graphs showing LVID;s (**A**), LVID;d (**B**), ESV (**C**), EDV (**D**), SV (**E**), EF (**F**), FS (**G**), and CO (**H**) a determined by M-mode echocardiography in 4-month-old male WT (n=8) and *Rbm20*^*ΔRS*^ (n=7) mice, as well as female WT (n=7) and *Rbm20*^*ΔRS*^ (n=11) mice. Data are shown as mean ± standard deviation. Dots represent measurements from individual animals. Two-way ANOVA with the Šídák method for multiple comparisons was performed to analyze the effect of sex and genotype on each of the aforementioned parameters. The interaction between sex and genotype was not significant for any parameter. ns, not significant; \**p*<0.05; \*\**p*<0.01; \*\*\**p*<0.001; \*\*\*\**p*<0.0001.

### Target gene splicing and expression are altered in the hearts of *Rbm20*^*ΔRS*^ mice

Disrupted splicing of RBM20 target genes is characteristic of RBM20 cardiomyopathy.^5-8^ To determine whether splicing of RBM20 targets is also disrupted in the hearts of *Rbm20*^*ΔRS*^ mice, RNA was extracted from the LVs of 2-month-old male WT and *Rbm20*^*ΔRS*^ mice, submitted for RNA-seq, and differentially spliced genes (DSGs) were identified. A total of 103 genes were differentially spliced in the hearts of *Rbm20*^*ΔRS*^ mice in comparison to WT controls, including several well-established RBM20 splicing targets such as *Ttn, Camk2d, Ryr2*, and *Tpm2* (Figure 3A and Table S4).^15,21^ To confirm disrupted splicing of *Ttn*, which is the primary splice target of RBM20,^15^ proteins were extracted from the ventricular myocardium of 2-month-old male *Rbm20*^*ΔRS*^ mice, and the expression of titin isoforms was assessed by agarose gel electrophoresis. Extracts prepared from age-matched male WT and *Rbm20*^*ΔRRM*^ mice served as negative and positive controls, respectively, for disrupted *Ttn* splicing.^17^ N2B titin was the predominant isoform expressed in the ventricle of age-matched WT mice (Figure 3B). On the other hand, titin was shifted to a giant N2BA (N2BA-G) isoform in *Rbm20*^*ΔRS*^ mice similar to that observed in *Rbm20*^*ΔRRM*^ mice (Figure 3B),^17^ confirming that *Ttn* splicing is impaired in the hearts of these mice. All five alternative splicing events were represented among the DSGs, including 65 events with skipped exons (SE), 6 with an alternative 5’ splice site (A5SS), 17 with an alternative 3’ splice site (A3SS), 12 with mutually exclusive exons (MXE), and 11 with retained introns (RI) (Figure 3C-D). The differential splicing of three other RBM20 target genes, *Camk2d, Ryr2*, and *Tpm2*, was also validated by RT-PCR and results were in line with the RNA-seq results (Figure S3).

**Figure 3.**
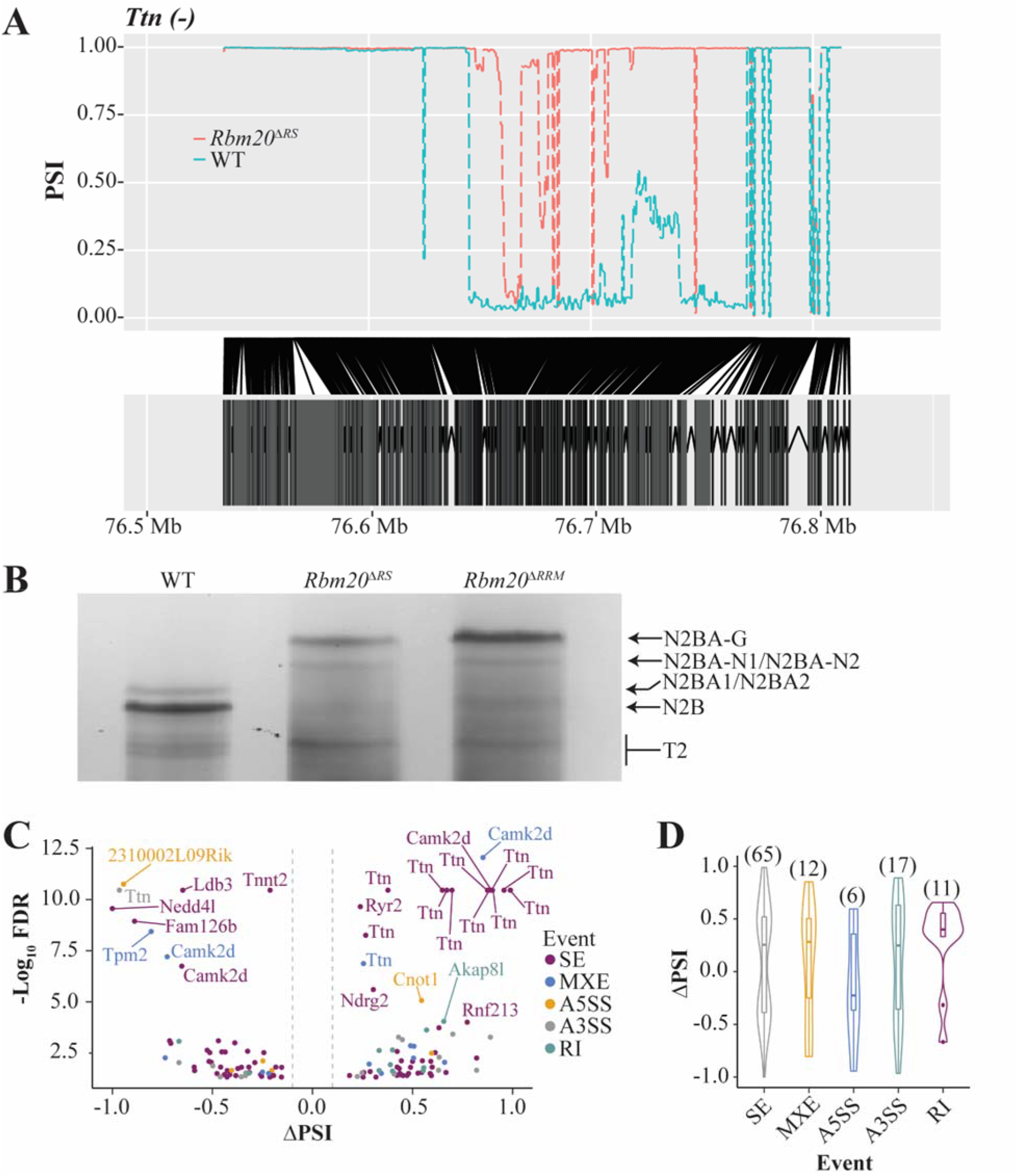
Splicing of RBM20 target genes is disrupted in the hearts of *Rbm20*^*ΔRS*^ mice. **A**, RNA-seq PSI alternative splicing maps for *Ttn* comparing WT (blue) and *Rbm20*^*ΔRS*^ (red). **B**, Titin isoforms detected in the LV myocardium of WT, *Rbm20*^*ΔRS*^, and *Rbm20*^*ΔRRM*^ mice. **C**, Volcano plot showing genes that are differentially spliced in the hearts of *Rbm20*^*ΔRS*^ mice relative to WT control. Genes with −;log_10_ FDR > 5 and ∣ΔPSI∣ > 0.1 are labeled. SE, skipped exon; MXE, mutually exclusive exons; A5SS, alternative 5′ splice site; A3SS, alternative 3′ splice site; RI, retained intron. **D**, Violin plots representing the distributions of statistically significant ΔPSI (percent spliced-in) values for the different classes of AS events in *Rbm20*^*ΔRS*^ mice relative to WT controls (ΔPSI = PSI(ΔRS) − PSI(WT)). The lower and upper bounds of the embedded box represented the 25^th^ and 75^th^ percentile of the distribution, respectively. The horizontal line in the box represented the median. The numbers of events are shown above each plot.

In addition to DSGs, 1399 genes were found to be differentially expressed in the hearts of *Rbm20*^*ΔRS*^ mice relative to that in age-matched WT controls (Figure S4 and Table S5). Of the 1399 genes, 595 and 824 were significantly up- and down-regulated, respectively (Figure S4 and Table S5). The most highly up-regulated gene was *Nppa*, a well-known cardiac stress marker (Figure S4 and Table S5). Also among the most highly up-regulated genes was another stress response protein *Ankrd1* (Figure S4 and Table S5), which has been shown to be up-regulated in heart failure,^22^ as well as DCM.^23^ In agreement with these changes, one of the most highly down-regulated genes was *Hopx* (Figure S4 and Table S5), which is down-regulated in heart failure in humans and mice.^24^ Collectively, these results further validate the development of cardiac stress and failure in *Rbm20*^*ΔRS*^ mice.

### RBM20 is mis-localized in the hearts of *Rbm20*^*ΔRS*^ mice

Recently, it was found that pathogenic mutations in the RBM20 RS domain promote nucleocytoplasmic shuttling and sarcoplasmic accumulation of the protein.^4-8^ Thus, we hypothesized that deletion of the RS domain in RBM20 prevents nuclear import and/or retention of the protein in the nucleus. To test this hypothesis, the localization of RBM20 was determined by IHC in cardiac sections from 2-month-old male WT and *Rbm20*^*ΔRS*^ mice. In age-matched male WT mice, RBM20 was localized to two discrete speckles in the nucleus previously shown to serve as sites for *Ttn* pre-mRNA processing (arrows, Figure 4A, right and Video S1).^25^ Consistent with our hypothesis, in *Rbm20*^*ΔRS*^ mice RBM20 was localized to perinuclear puncta resembling the RBM20 granules previously reported in humans and other animals carrying RBM20 missense mutations (arrows, Figure 4B, right and Video S2).^4-8^ The localization of RBM20 was also assessed in the hearts of 2-month-old male *Rbm20*^*ΔRRM*^ mice. In contrast to RBM20 lacking the RS domain, RBM20 with the RRM was localized exclusively within the nucleus (bracket, Figure 4C, right). Interestingly, in *Rbm20*^*ΔRRM*^ mice, RBM20 did not localize to nuclear speckles, but instead exhibited a diffuse nuclear localization pattern (Figure 4C and Video S3).

**Figure 4.**
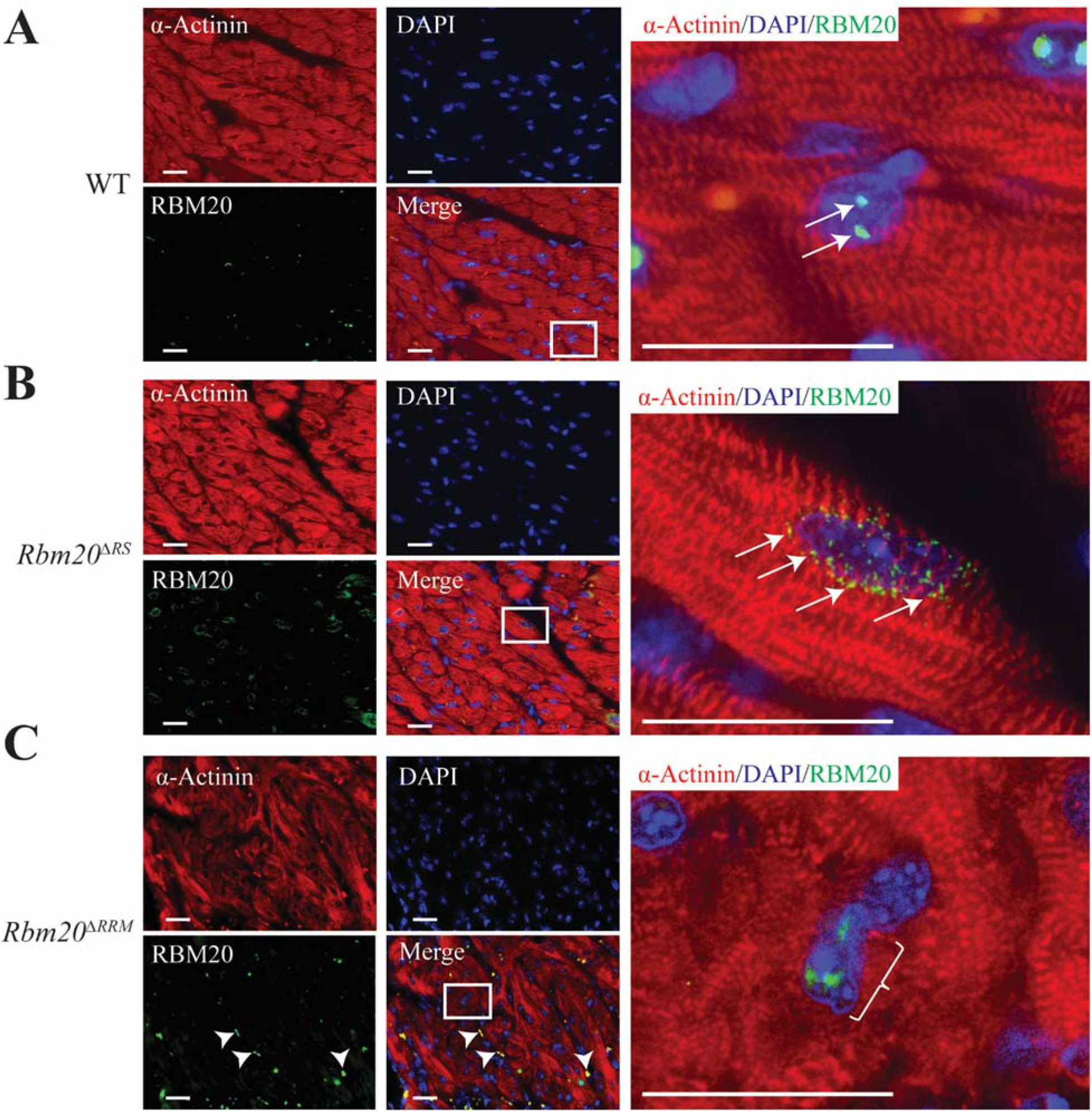
Loss of the RS, but not RRM, domain in RBM20 promotes sarcoplasmic localization. **A-C**, Representative IHC images showing the localization of RBM20 in the hearts of WT (**A**), *Rbm20*^*ΔRS*^ (**B**), and *Rbm20*^*ΔRRM*^ (**C**) mice. Inset images are shown on the right. Arrows and backet denote RBM20 staining. Arrow heads mark non-specific staining of non-cardiomyocytes. Scale bars are 25 μm.

### The RS domain of RBM20 mediates nuclear localization

To verify that the RS domain mediates RBM20 nuclear localization, we generated rat RBM20 constructs containing WT RBM20, RBM20 lacking the entire region from the start of the RRM domain to the end of the RS domain (Δ522-658), RBM20 lacking the RRM domain (ΔRRM), RBM20 lacking a classical monopartite NLS (KRYK) located between the RRM and RS domains that has been described previously (ΔRYK-KK),^26^ and RBM20 lacking the RS domain (ΔRS) (Figure 5A).

**Figure 5.**
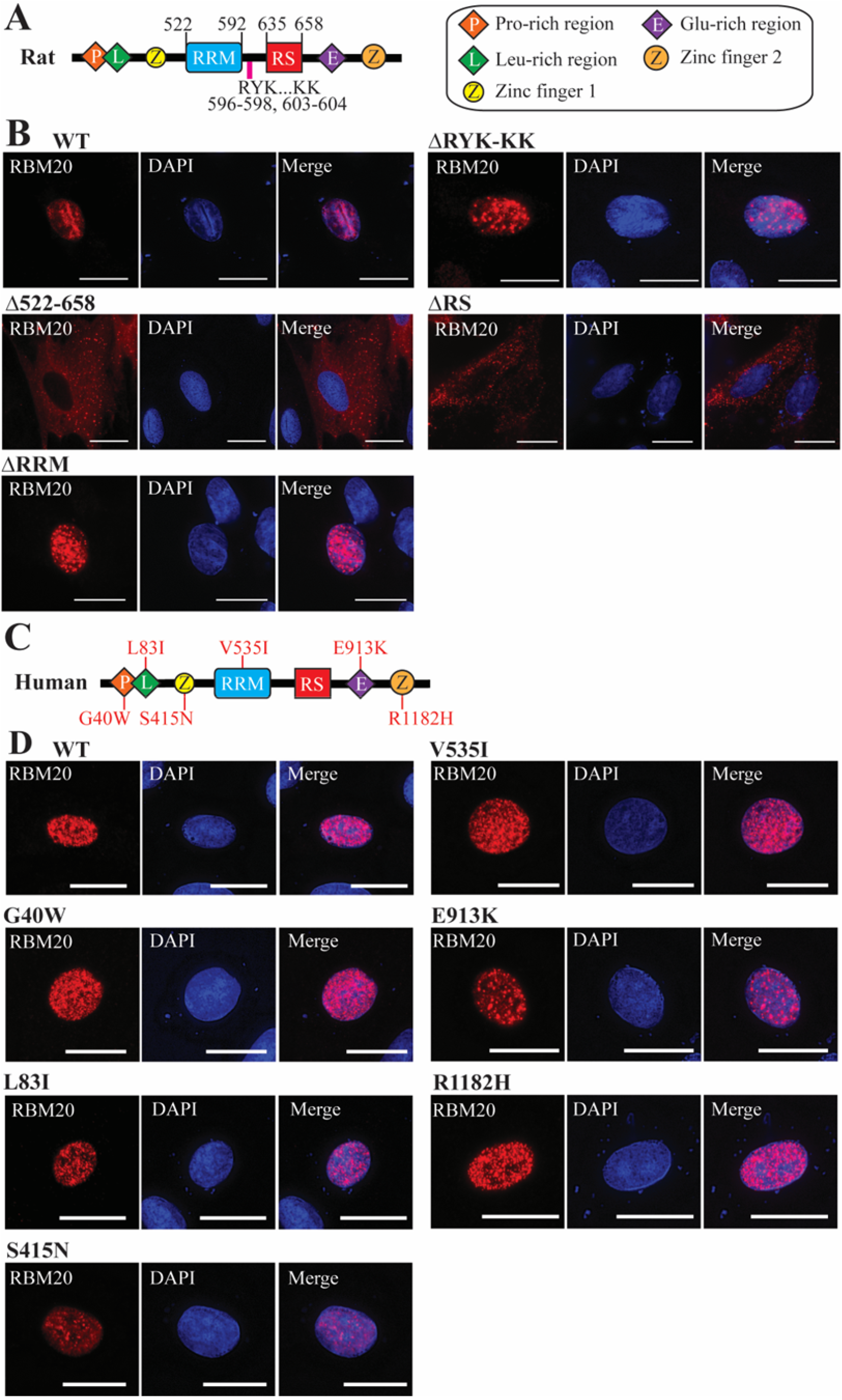
The RS, but not the RRM, domain controls RBM20 nuclear localization. **A**, Schematic showing domain structure of rat RBM20. **B**, Representative ICC images showing rat RBM20 domain and sequence deletion construct localization in transfected H9c2 cells. Scale bars are 20 μm. **C**, Schematic showing domain structure of human RBM20 with DCM-associated mutations in the respective domains/regions within the protein listed. **D**, Representative ICC images showing the localization of human RBM20 constructs harboring DCM-associated mutations in domains/regions other than the RS domain expressed in H9c2 cells. Scale bars are 20 μm.

To establish the role of these domains and sequence elements in mediating RBM20 nuclear localization, H9c2 cells were transfected with the different constructs and the localization of RBM20 was established by ICC. As expected, WT RBM20 was exclusively localized within the nucleus of transfected cells (Figure 5B). Conversely, the Δ522-658 construct localized to the cytoplasm of transfected H9c2 cells (Figure 5B).^27^ Unlike the Δ522-658 construct, both the ΔRRM and ΔRYK-KK constructs were contained within the nucleus (Figure 5B), indicating that these sequences are not necessary for nuclear localization. Localization of the ΔRRM construct to the nucleus of transfected H9c2 was in agreement with the localization of RBM20 in the hearts of *Rbm20*^*ΔRRM*^ mice (Figure 4C). In contrast, the ΔRS construct localized to the cytoplasm in transfected cells (Figure 5B), which is consistent with the sarcoplasmic accumulation of RBM20 observed in *Rbm20*^*ΔRS*^ mouse hearts (Figure 4B).

The RS domain of RBM20 is one of three mutation hotspots in the protein with the other two being the RRM and E-rich region.^19,20^ Notably, a growing list of mutations in other domains/regions of the protein have also been reported in the context of DCM. To confirm that DCM-associated mutations in other domains/regions outside the RS domain do not impair nuclear localization of the protein, we engineered the G40W,^28^ L83I,^29^ S415N, V535I,^30^ E913K,^31^ and R1182H^28,32^ mutations into the respective domains/regions of human RBM20 and the localization of these constructs was determined by ICC in transfected H9c2 cells (Figure 5C). It should be noted that although the S415N mutation has not been reported in the literature, according to information in the ClinVar database (ClinVar accession: VCV000548140.4) this variant was discovered in a female patient with early onset dilated cardiomyopathy and heart failure. Consistent with the dominant role played by the RS domain in determining RBM20 nuclear localization, all mutant constructs were confined to the nucleus of transfected cells (Figure 5D). Collectively, these results indicate that only the RS domain of RBM20 is responsible for mediating nuclear localization of the protein.

### The D1 sequence element constitutes the core NLS in RBM20

To elucidate the mechanism underlying RS domain-mediated control of RBM20’s nuclear localization, *in silico* analysis was used to identify putative NLS sequence elements within the RBM20 RS domain using the core transportin-interacting SRSF1 RS domain sequence (i.e., RSRSRSRSR)^33^ as a template. The core transportin-interacting SRSF1 RS domain sequence was chosen as a template because SRSF1 is the prototypical SR protein and interactions between this sequence and the nuclear import receptor *Tnpo3* have been studied previously.^33^ *In silico* analysis revealed the presence of two core SRSF1 RS domain sequence-like elements located within the RS domain of RBM20 hereafter referred to as D1 and D2 (Figure 6A). To determine whether one or both sequence elements are required for RBM20 nuclear localization, we made rat RBM20 constructs lacking both D1 and D2 (ΔD12), D1 only (ΔD1), D2 only (ΔD2), D3 only (ΔD3), and both D2 and D3 (ΔD23). These constructs were transfected in H9c2 cells, and the localization of RBM20 was determined by ICC. Like the ΔD12 construct, the ΔD1 construct localized exclusively to the cytoplasm of H9c2 (Figure 6B). In contrast, the ΔD2, ΔD3, and ΔD23 constructs were all localized within the nucleus (Figure 6B).

**Figure 6.**
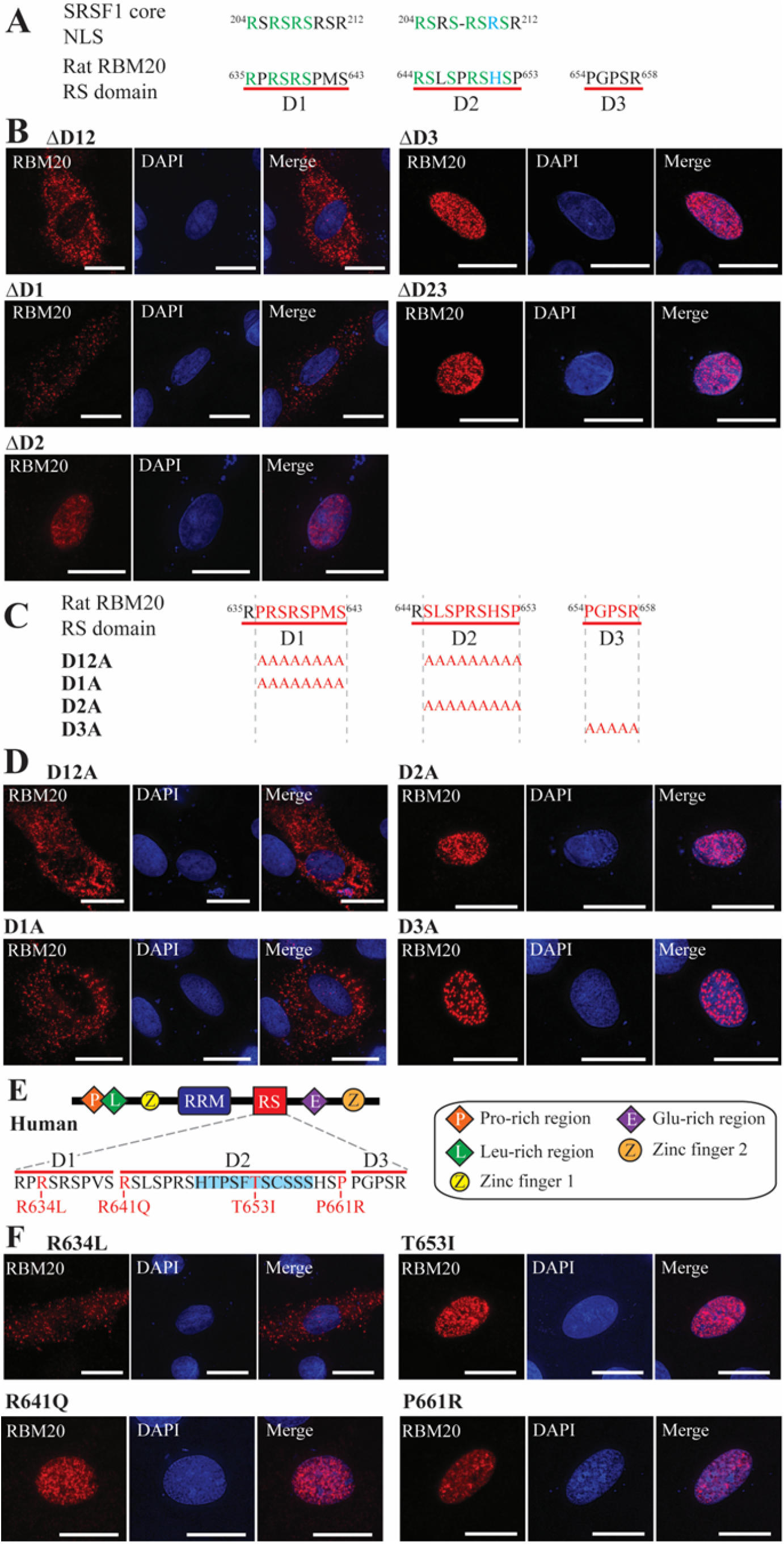
Disruption of the D1 core NLS sequence element abolishes RBM20 nuclear localization. **A**, *In silico* identification of SRSF1 core NLS-like sequence elements in the RS domain of rat RBM20. Conserved amino acids are shown in green and blue denotes a positively charged amino acid. **B**, Representative ICC images showing rat RBM20 deletion construct localization in transfected H9c2 cells. **C**, Schematic showing rat RBM20 Ala substitution constructs for *in vitro* expression in H9c2 cells. **D**, Representative ICC images showing rat RBM20 Ala substitution construct localization in transfected H9c2 cells. Scale bars are 20 μm. **E**, Schematic showing human RBM20 domain structure with a zoom in on the RS domain sequence and DCM-associated mutations listed. The human-specific segment in the D2 sequence element is highlighted in light blue. **F**, Representative ICC images showing the localization of human RBM20 constructs harboring DCM-associated mutations in the D1 and D2 sequence elements. Scale bars are 20 μm.

Rat RBM20 mutant constructs with Ala residues in place of amino acids 636-643 in D1, 645-653 in D2, or 654-658 in D3 were generated to further validate the necessity of the D1 sequence element for nuclear import. The D12A construct contained Ala substitution for both D1 and D2 (Figure 6C). In agreement with the results of the previous deletion experiments, constructs in which the D1 sequence element was replaced with Ala residues (D1A and D12A) exhibited a cytoplasmic localization pattern while those containing an intact D1 sequence element (D2A and D3A) were located within the nucleus (Figure 6D).

In addition to the growing list of mutations reported in the D1 sequence element, several DCM-associated mutations have also been reported in D2, including variants in a human-specific stretch within this sequence element (Figure 6E). To determine whether DCM-associated mutations in the human D2 sequence element promote RBM20 re-localization, the R641Q,^28^ T653I,^34^ and P661R (ClinVar accession: VCV000965264.3) mutations were engineered into human RBM20, and the mutant constructs were expressed in H9c2 cells. The R634L mutation,^35^ which is located in the D1 sequence element, served as a positive control for disrupted nuclear localization. In agreement with the D1/D2 deletion and Ala replacement experiments in rat RBM20 (Figure 6A-D), only the R634L mutation promoted accumulation of human RBM20 in the cytoplasm of transfected H9c2 cells (Figure 6F). Collectively, these results confirm that the D1 sequence element is the core NLS in RBM20 and suggest that mutations in the D1 core NLS alone promote RBM20 cardiomyopathy via impaired nuclear import and cytoplasmic accumulation of the protein.

### Pseudo-phosphorylation of the RS domain of RBM20 does not rescue nuclear localization

Recently, it was found that multiple Ser residues in the RS domain of RBM20, including all three Ser residues in the D1 sequence element, are basally phosphorylated.^8,36^ Given that RS domain phosphorylation has previously been shown to be important for nucleocytoplasmic shuttling of SR proteins such as SRSF1,^37,38^ we next sought to establish the role of RS domain phosphorylation in RBM20 nuclear localization. Constructs were generated in which all phosphorylatable Ser residues within the RS domain were mutated to either non-phosphorylatable Ala (S8A) residues or phosphomimetic Asp residues (S8D) (Figure 7A). Mutation to Ser residues prevented RBM20 nuclear import (Figure 7B). Conversely, mutation to the phosphomimetic residue Asp did not rescue nuclear localization of RBM20, suggesting that phosphorylation is not necessary for nuclear import of the protein (Figure 7B).

**Figure 7.**
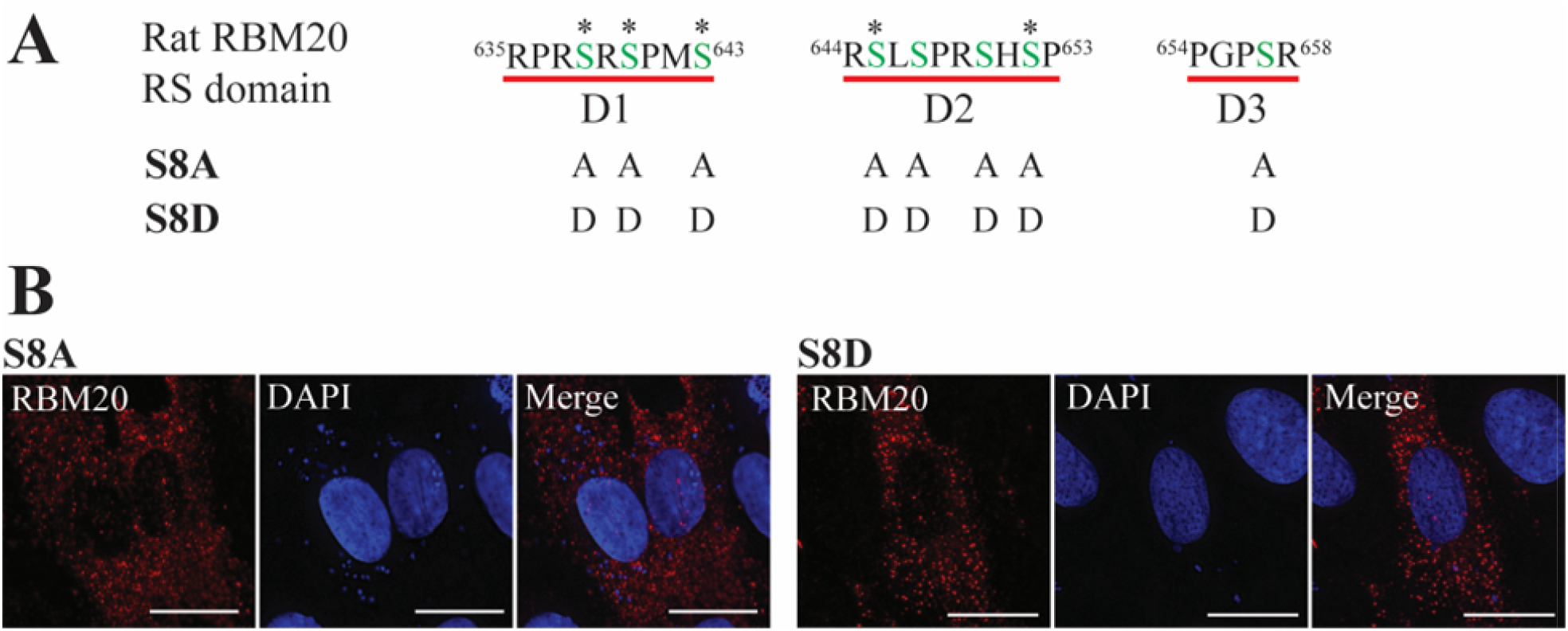
Replacement of phosphorylatable Ser residues in the RBM20 RS domain with phosphomimetic Asp residues does not rescue nuclear localization. **A**, Schematic showing rat RBM20 Ala and Asp mutation constructs for *in vitro* expression in H9c2 cells. Asterisks denote Ser residues previously identified as phosphorylation sites.^8,36^ **B**, Representative ICC image showing RBM20 Ala and Asp mutation construct localization in transfected H9c2 cells. Scale bars are 20 μm.

### D1 sequence integrity is critical for RBM20 nuclear localization

Prior studies have shown that the DCM-associated S635A^5,8^, R636S^7,27^, S637G^6^, and P638L^4,8^ mutations in RBM20, which are located in the D1 sequence element, disrupt nuclear import of the protein. To further assess the role that the amino acid composition of the D1 sequence element plays in mediating RBM20 nuclear localization, we examined the localization of RBM20 harboring five additional DCM-associated mutations in the D1 NLS for which no available localization data has been reported. The selected mutations included R634L, R634Q, R634W, R636C, and R636H (Figure 8A). The R636S mutation served as a positive control for disrupted nuclear localization (Figure 8A).^7,27^ The mutations were engineered into rat RBM20, and the subcellular localization of the mutant proteins was assessed following transfection in H9c2 cells using ICC. Consistent with previous reports,^7,27^ RBM20 harboring the R636S mutation (R639S in rat) exhibited a near exclusive cytoplasmic localization pattern (Figure 8B). In agreement with the necessity of D1 sequence integrity, all five of the selected mutations also disrupted the nuclear import of RBM20 and were localized to the cytoplasm (Figure 8B). To better understand the severity of these amino acid substitutions, Grantham’s distances (d), which provide a proxy for the interchangeability of two amino acids based on their composition, polarity, and molecular volume,^39^ were employed. While it is perhaps not surprising that the R636C mutation (d=180; both residues are polar, but Cys in uncharged and is smaller in terms of molecular volume) impacts RBM20 nuclear localization as this represents a radical substitution based on the classification system established by Li et al.,^40^ it is surprising that even the conservative amino acid substitutions R634Q (d=43; both residues are polar although Arg is charged while Gln is uncharged) and R636H (d=29; both residues are polar and basic differing primarily in molecular volume with His being smaller than Arg) were sufficient to disrupt RBM20 nuclear localization (Figure 8C).

**Figure 8.**
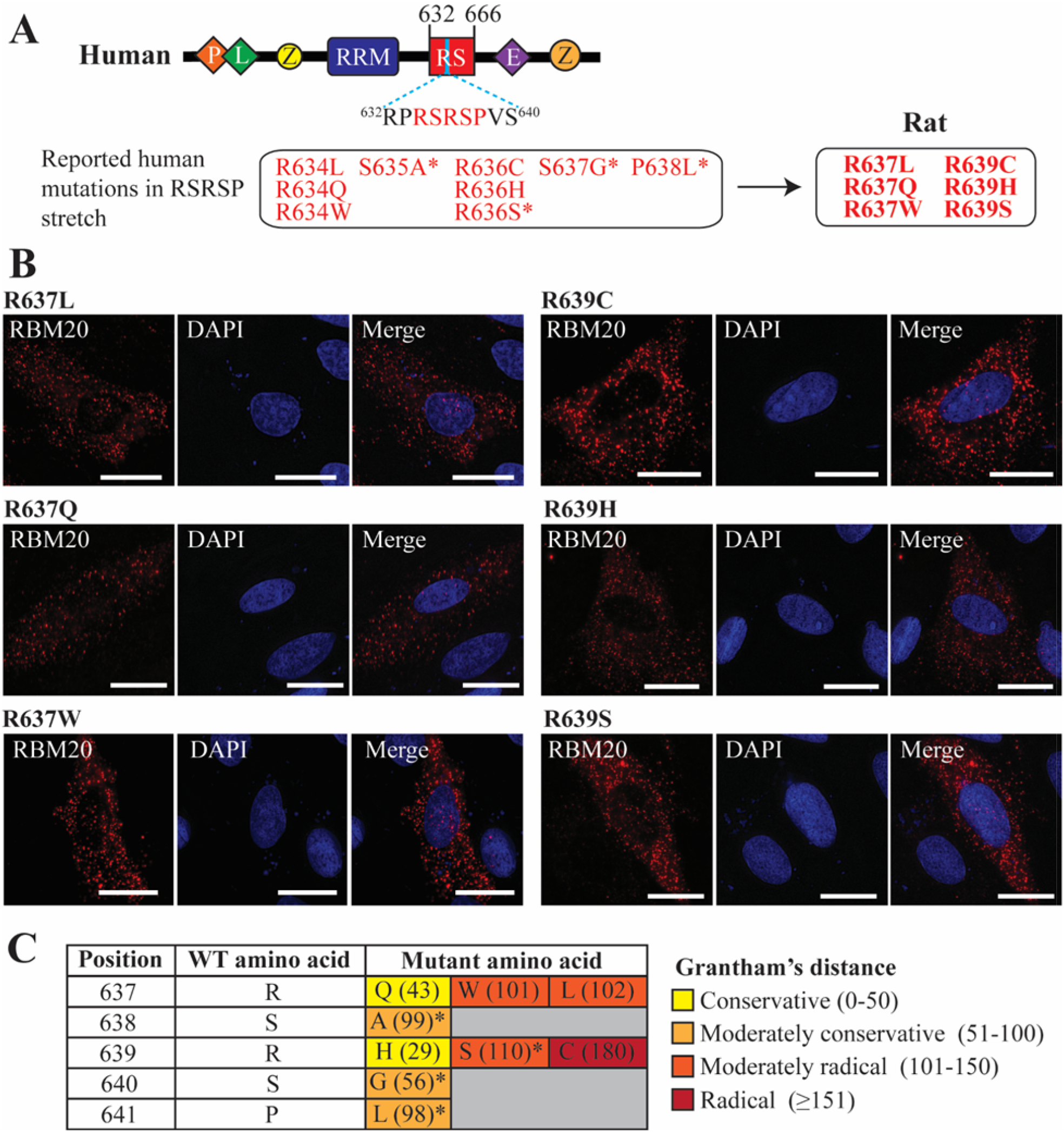
DCM-linked mutations in the RSRSP stretch contained within the D1 sequence element disrupt RBM20 nuclear localization. **A**, Schematic showing the domain structure of human RBM20 with the D1 sequence element shown. Critical RSRSP stretch is highlighted in red. Reported mutations in the RSRSP stretch in human RBM20 are listed along with the corresponding mutations in rat RBM20. Asterisks denote RBM20 mutants with localization data already available in the literature. **B**, Representative images showing the localization of rat RBM20 harboring DCM-linked mutations corresponding those reported in the RSRSP stretch of human RBM20. Scale bars are 20 μm. **C**, Table displaying Grantham’s distances for DCM-linked amino acid mutations in the RSRSP stretch that disrupt RBM20 nuclear localization. Amino acid positions are given for rat RBM20. The number in parentheses corresponds to Grantham’s distance between the amino acid in the WT sequence and the mutant amino acid. Grantham’s distances were classified as conservative, moderately conservative, moderately radical, or radical as proposed by Li et al.^40^ Asterisks denote RBM20 mutants with localization data already available in the literature.

## DISCUSSION

There are three primary mechanisms contributing to RBM20 cardiomyopathy. These are 1) a reduction in titin-based passive stiffness resulting from changes in titin isoform expression,^15,17,41^ 2) impaired contractile function secondary to altered splicing of Ca^2+^-handling genes,^16^ and 3) pathophysiological gain-of-function effects caused by RBM20 nucleocytoplasmic shuttling and sarcoplasmic accumulation in RBM20 granules.^4-8^ However, it remains unclear which of these mechanisms is the driving force behind the development of DCM. Notably, multiple studies have now shown that mice with disrupted splicing of RBM20 target transcripts do not develop an overt DCM phenotype,^17,18^ indicating that altered splicing alone does not significantly contribute to the development of RBM20 cardiomyopathy. Herein, employing a novel RBM20 RS domain deletion mouse model in combination with *in vitro* experiments, we show that loss of RBM20 nuclear localization, rather than disrupted splicing, is the driving mechanism in DCM. Based on this finding, we assert that RBM20 nucleocytoplasmic trafficking and accumulation in cytoplasmic RBM20 granules underlies the development of RBM20 cardiomyopathy, although disruption of an unknown non-splicing nuclear function of RBM20 cannot be ruled out at this time. This claim is supported by the fact that animals carrying RBM20 mutations that promote RBM20 granule formation in the sarcoplasm develop severe DCM.^5-8^ With the shift in mechanistic focus from mis-splicing to RBM20 granules as the causative mechanism in RBM20-related cardiomyopathy, an important next step will be to determine whether preventing nucleocytoplasmic trafficking of RBM20 by stabilizing the complex formed between importer proteins and the disrupted core NLS sequence or targeted disassembly of RBM20-mediated granules can benefit RBM20 cardiomyopathy.

One question that remains unanswered is why *Rbm20*^*ΔRS*^ mice did not exhibit the early mortality previously observed in mice harboring pathogenic RS domain mutations^6,8^ even though RBM20 granules appeared to be present. We believe that this may be because the RS domain plays an important role in determining disease severity by influencing the composition of RBM20 granules. Indeed, the RS domain in SR proteins is known to mediate a host of protein-protein interactions, including those essential for assembly of the spliceosome.^14^ Our present hypothesis is that residual RS domain function in the context of pathogenic mutations in RBM20 enables the recruitment of a specific complement of proteins and/or RNAs to RBM20 granules, leading to a more severe DCM phenotype. Nevertheless, additional studies will be necessary to validate this hypothesis.

Another important finding of the present study is that the RS domain, not the RRM, controls RBM20 nuclear localization. The findings of a previous study conducted by Filippello et al. suggested that additional sequences in RBM20, specifically the RRM and sequence between the RRM and RS domains, may be required for nuclear retention of the protein.^26^ Herein, we provide two lines of evidence indicating that the RS domain is solely responsible for RBM20 nuclear localization. First, RBM20 nuclear localization is disrupted in the hearts of *Rbm20*^*ΔRS*^ mice. Second, follow up experiments in transfected H9c2 cells demonstrated that, while the RRM was dispensable for nuclear localization, the RS domain was necessary. Consistently, DCM-associated mutations in domains other than the RS domain failed to re-localize RBM20 to the cytoplasm in transfected cells. The latter results are in agreement with the findings of previous *in vitro* experiments indicating that DCM-associated mutations outside the RS domain do not promote RBM20 re-localization.^36^

Consistent with the role for the RS domain in controlling RBM20 nuclear import, we identified the D1 sequence element as the core NLS in RBM20. This finding is supported by the observation that mutations in the RSRSP stretch, which is located within the D1 core NLS, promote nucleocytoplasmic transport and accumulation of the protein in the sarcoplasm.^5-8^ Surprisingly, while not all possible amino acid substitution permutations have been tested, here we show that even conservative amino acid substitutions, such as R634Q and R634H, within this sequence are sufficient to disrupt nuclear localization of the protein. Thus, the RS domain and, more specifically, the integrity of the D1 core NLS is essential for nuclear import of RBM20. Identification of the nuclear import receptor for RBM20 will be a key step for understanding how mutations in this NLS disrupt nuclear import.

Prior studies have shown that phosphorylation of the RS domain in SRSF1, the prototypical SR protein, is also important for interactions with SR-specific transportin proteins and, thus, for nuclear import of the protein.^33,37,38^ In the case of RBM20, Murayama et al. demonstrated that mutation of two phosphorylated Ser residues (S637 and S639 in mice, or S635 and S637 in humans) in the RS domain of RBM20 disrupted splicing of a *Ttn* reporter and promoted nuclear exclusion of the protein.^36^ We have reported similar findings; yet, replacement of these two Ser residues with the phosphomimetic amino acid Asp, which has previously been shown to at least partially replace phosphorylated Ser with respect to mediating SR protein nuclear import,^42^ did not rescue nuclear localization.^8^ Importantly, we recently identified several additional sites of phosphorylation within the RS domain using middle-down mass spectrometry,^8^ raising the question of whether phosphorylation at other sites regulates RBM20 nuclear localization. In this study, we mutated all phosphorylatable Ser residues in the RS domain of rat RBM20 to either Ala or Asp residues. Surprisingly, replacement of Ser residues in the RS domain with phosphomimetic Asp residues did not restore nuclear localization of RBM20. Taken together, these results suggest that phosphorylation of the RBM20 RS domain is not required for nuclear localization. Regardless, a notable caveat is that, to date, all studies that have examined the role of RBM20 phosphorylation in nuclear localization have employed mutagenesis to prevent phosphorylation at specific sites in the protein.^8,36^ Given that mutations themselves disrupt RBM20 localization, assessing the role phosphorylation plays in subcellular localization by methods other than mutagenesis will be a necessary future step for confirming these findings. Moreover, these experiments cannot exclude the possibility that phosphorylation at specific sites or combinations of sites and dephosphorylation at others is necessary for nuclear import.

## METHODS

### Experimental animals and tissue collection

*Rbm20*^*ΔRS*^ mice were generated via CRISPR/Cas9 genome editing in collaboration with the Advanced Genome Editing Animal Models Core Facility at the University of Wisconsin-Madison by introducing Cas9 protein (6 μM, IDT v3), two sgRNAs (3.3 μM each, IDT sgRNAs), and a single-stranded donor (20 ng/μl) into C57BL/6J embryos by electroporation (sequence information for single-stranded donor and sgRNAs are provided in Table S1). Founders were screened using Illumina targeted deep sequencing to identify animals carrying perfect excisions of the region between E633-S666 in RBM20. Founders were backcrossed to C57BL/6J mice to generate F1s, which were similarly characterized by targeted deep sequencing. Heterozygous founders were crossed to produce homozygous *Rbm20*^*ΔRS*^ mice for subsequent characterization.

For genotyping, genomic DNA was isolated from toe clips by incubating in DirectPCR (Tail) Lysis Reagent for Genotyping Crude Lysates (cat# 102-T; Viagen Biotech, Los Angeles, CA, USA) with proteinase K (cat# DSP50220-0.1; Dot Scientific, Burton, MI, USA) for 12 hr at 55 °C followed by a 45 min heat inactivation at 85 °C. Isolated genomic DNA was combined with EconoTaq PLUS GREEN 2X Master Mix (cat# 30033-1; Lucigen, Middleton, WI, USA) and forward and reverse primers (primer information listed in Table S2) and PCR was carried out with an initial denaturation at 94°C for 2 min, followed by 35 cycles of 94 °C for 15 sec, 63 °C for 30 sec, and 72 °C for 30 sec. *Rbm20*^*ΔRRM*^ mice are also in the C57BL/6J background and have been described previously.^17^ WT C57BL/6J mice served as controls for all experiments.

All mice were maintained on standard rodent chow (cat# 8604; Envigo, Indianapolis, IN, USA) and were housed in a facility on a 12 hr/12 hr light/dark cycle with access to food and water ad libitum. Hearts were collected from 2- and 4-month-old mice immediately following euthanasia. After removal, hearts were either paraffin-embedded for histology and/or immunohistochemistry (IHC) or sectioned into chambers, snap-frozen in liquid nitrogen, and stored in a −80 °C freezer for later biochemical experiments. All procedures involving animals were carried out following the recommendations in the Guide for the Care and Use of Laboratory Animals published by the National Institutes of Health and were approved by the Institutional Animal Care and Use Committee of the University of Wisconsin-Madison.

### Histology

Whole hearts from 2- or 4-month-old WT, *Rbm20*^*ΔRS*^, and *Rbm20*^*ΔRRM*^ mice were isolated and fixed with 4% paraformaldehyde, paraffin-embedded, sectioned, and stained with Masson’s trichrome stain. Stained sections were photographed using a Keyence BZ-X800 All-in-one Fluorescence Microscope (Itasca, IL, USA), and the fibrotic area was quantified using ImageJ.^43^ The proportion of fibrotic area was calculated as a ratio of the fibrotic area to the total area.

### Echocardiography

Transthoracic echocardiography was performed using a Visual Sonics Vevo 3100 ultrasonograph (Toronto, ON, Canada) outfitted with a MX400 transducer, ∼30-MHz, as detailed previously.^44^ For acquisition of two-dimensional guided M-mode images at the tips of papillary muscles and Doppler studies, mice were sedated by facemask administration of 1% isoflurane, hair was removed, and mice were maintained on a heated platform.

End diastolic and systolic LV diameter, as well as anterior and posterior wall (AW and PW respectively) thickness, were measured on-line from M-mode images obtained in a parasternal long axis view using the leading edge-to-leading edge convention. All parameters were measured over at least three consecutive cardiac cycles and averaged. LV fractional shortening (FS) was calculated as [(LV diameter_diastole_ – LV diameter_systole_)/LV diameter_diastole_] x 100; ejection fraction (EF) as [(7.0/(2.4 + LV diameter_diastole_)(LV diameter_diastole_)^3^ - (7.0/(2.4 + LV diameter_systole_)(LV diameter_systole_)^3^/(7.0/(2.4 + LV diameter_diastole_)(LV diameter_diastole_)^3^ x 100; and LV mass was calculated by using the formula [1.05 x ((Posterior Wall_diastole_ + Anterior Wall_diastole_+ LV diameter_diastole_)^3^ – (LV diameter_diastole_)^3^)]. Heart rate was determined from at least three consecutive intervals from the pulse wave Doppler tracings of the LV outflow tract. The same person obtained all images and measures.

### IHC

Paraffin-embedded sections of heart were deparaffined and rehydrated by washing twice with pure xylene, once with 50:50 xylene:ethanol, twice with pure ethanol, and then once each with 90%, 80%, and 70% ethanol in water, followed by a final wash with Milli-Q water. Subsequently, sections were soaked in 0.5% Triton X-100 (cat# X100; Sigma-Aldrich, St. Louis, MO, USA) for 10 min to permeabilize and then rinsed three times with Milli-Q water. Antigen retrieval was performed by soaking in Tris-EDTA buffer (10 mM Tris, 1 mM EDTA, 0.05% Tween 20, pH 9.0) and boiling at approximately 98 °C for 20 min. Slides were cooled to RT, blocked by incubating in blocking buffer [PBS containing 5% goat serum (cat# G6767; Sigma-Aldrich), 0.1% Triton X-100 (cat# X100; Sigma-Aldrich), and 0.05% Tween 20 (cat# P20370-0.5; Research Products International, Mount Prospect, IL, USA)] at RT for 1 hr, and then incubated with primary antibodies in blocking buffer [1:400 homemade rabbit anti-RBM20,^15^ or 1:500 anti-α-actinin (cat# A7811; Sigma-Aldrich)] overnight at 4 °C. Next, slides were washed with TBST and incubated with secondary antibodies in blocking buffer [goat anti-rabbit (cat# A32731; Invitrogen, Waltham, MA, USA) or goat anti-mouse (cat# 8890; Cell Signaling Technology, Danvers, MA, USA)] for 1 hr at RT, washed with TBST, and mounted in SlowFade™ Gold Antifade Mountant with DAPI (cat# S36938; Invitrogen). Slides were photographed using a BZ-X800 All-in-one Fluorescence Microscope (Keyence).

### Titin gel electrophoresis

Titin isoforms were resolved as previously described using a 1% vertical sodium dodecyl sulfate (SDS)-agarose gel electrophoresis (VAGE) system.^45,46^ Frozen LV myocardium from 2-month-old mice was homogenized in urea-thiourea-SDS-dithiothreitol sample buffer using a 2010 Geno/Grinder (SPEX SamplePrep, Metuchen, NJ, USA) with 5 cycles of 1500 strokes/min for 1 min and 15 sec rest between cycles (repeated four times for a total of 20 cycles) and subsequently incubated at 55 °C for 10 min. The denatured protein samples were loaded on a 1% SDS-agarose gel and run at a constant current of 30 mA for 3.5 hr. The agarose gel was fixed by incubating in fixing solution (50% methanol, 12% glacial acetic acid, and 5% w/v glycerol) for 1 hr at RT and then dried overnight at 37 °C. The dried gel was silver stained as previously reported^45,46^ and imaged using a ChemiDoc Imaging System (Bio-Rad, Hercules, CA, USA).

### RNA preparation and RNA-sequencing (RNA-seq)

Total RNA was extracted from the LV myocardium of 2-month-old male WT and *Rbm20*^*ΔRS*^ mice (n = 3 per group) using TRIzol Reagent (Life Technologies, Waltham, MA, USA) and treated with DNase I (cat# D9905K; Lucigen, Middleton, WI, USA). The concentration and purity of the isolated RNA were determined using a NanoDrop One Microvolume UV-Vis Spectrophotometer (Thermo Fisher Scientific, Waltham, MA, USA) and electrophoresis prior to submission for RNA-seq. RNA-seq was performed by the University of Wisconsin-Madison Biotechnology Center Gene Expression Center & DNA Sequencing Facility. Quality check of the raw data was performed using FastQC software.^47^ Low quality reads and adapter sequences were trimmed using Trimmomatic.^48^ Trimmed reads were mapped to the mouse reference genome (Mus musculus GRCm39) using STAR.^49^

### Alternative splicing analysis

Analysis of differentially spliced genes in the hearts of *Rbm20*^*ΔRS*^ mice relative to WT (control) was performed in Multivariate Analysis of Transcript Splicing software for replicates (rMATS).^50^ Five types of alternative splicing events were analyzed, including skipped exon (SE), alternative 5’ splice site (A5SS), alternative 3’ splice site (A3SS), mutually exclusive exons (MXE), and retained intron (RI). The output from rMATS was filtered using false discovery rate (FDR) ≤ 5% and an absolute value of ΔPSI ≥ 10% as the cut-off criteria to identify significant alternative splicing events. Volcano plots and violin plots were plotted using the “ggplot2” software package (v3.3.4).^51^

### PSI analysis

Sequences from RNA-seq analysis were trimmed with Trimmomatic^48^ with parameters “SLIDINGWINDOW:4:15 MINLEN:36”. The trimmed sequences were then mapped to GRCm39 reference genome from Ensembl release 107 with STAR.^49^ PSI data were then calculated with DEXSeq^52^ and PSI.sh^53^ with modification adapted to Python3 and new output format of STAR. The output result was visualized using ggbio.^54^

### Differentially expressed genes analysis

For gene-level expression, gene counts were estimated using the “--quantMode GeneCounts” option in STAR.^49^ The R package edgeR^55^ was used to normalize gene counting based on trimmed mean of M-values method.^56^ Only expressed genes with at least 15 counts in at least three samples were evaluated, resulting in 14,787 genes for further analysis. Analysis of differentially expressed genes was carried out for pairwise comparisons between *Rbm20*^*ΔRS*^ and WT based on a negative binomial generalized linear model using the edgeR package.^55^ The statistical tests were corrected for multiple testing and only genes with a FDR less than 0.05 were considered significant.^57^ Enrichment network analysis was performed using Metascape^58^ (http://metascape.org) under default model. Volcano plots and heatmap were plotted using the “ggplot2” software package (v3.3.4).

### RT-PCR for alternative splicing events validation

The conditions used for the synthesis of cDNA and RT-PCR analysis have been reported previously.^8^ Briefly, cDNA was synthesized using iScript Reverse Transcription Supermix (cat# 1708841; Bio-Rad) following the manufacturer’s instructions. RT-PCR was carried out with EconoTaq PLUS GREEN 2X Master Mix (cat# 30033-1; Lucigen) and an initial denaturation at 94°C for 2 min, followed by 35 cycles of 94 °C for 15 sec, annealing for 30 sec (see Table S2 for primer information and anneal temperatures), and 72 °C for 30 sec. The housekeeping gene *Gapdh* was amplified by 25 cycles. PCR products were resolved on agarose gels and imaged using a ChemiDoc Imaging System (Bio-Rad).

### Plasmids

The rat pEGFP-C1-RBM20 WT mutation and deletion vectors were based on rat *Rbm20* CDS sequence (NM_001107611.2) and were constructed by General Biosystems, Inc. (Durham, NC, USA). The human pEGFP-C1-RBM20 WT and mutation vectors were based on human *RBM20* CDS sequence (NM_001134363.3) and were constructed by Gene Universal Inc. (Newark, DE, USA). An 8xHis-tag was added at the C-terminus of RBM20.

### Cell culture, transfection, and immunocytochemistry (ICC)

H9c2 cells were maintained at 5% CO_2_ and 37 °C in HyClone Dulbecco’s Modified Eagle Medium (DMEM) with high glucose (cat# SH30022.01; Cytiva, Marlborough, MA, USA) supplemented with 10 % fetal bovine serum (cat# SH30910.03HI; Cytivia), 1% Penicillin/Streptomycin (cat# SV30010; Cytiva), and 1% Sodium Pyruvate (cat# 11360070; Gibco, Thermo Fisher Scientific). Transfection was performed using FuGENE HD Transfection Reagent (cat# E2311; Promega, Madison, WI, USA) in accordance with the manufacturer’s instructions.

For ICC, H9c2 cells were grown on glass coverslips. Forty-eight hours post-transfection, cells were fixed with methanol for 15 min on ice. Following fixation, cells were blocked/permeabilized with PBS containing 5% goat serum (cat# G6767; Sigma-Aldrich), 0.1% Triton X-100 (cat# X100; Sigma-Aldrich), and 0.05% Tween 20 (cat# P20370-0.5; Research Products International) for 1 hr at RT. Subsequently, cells were incubated with homemade anti-RBM20 (1:400) primary antibody in blocking buffer [PBS containing 5% goat serum (cat# G6767; Sigma-Aldrich), 0.1% Triton X-100 (cat# X100; Sigma-Aldrich), and 0.05% Tween 20 (cat# P20370-0.5; Research Products International)] overnight at 4°C. After washing, cells were incubated with Alexa fluor-conjugated secondary antibodies (1:1500; cat# A11036 or cat# A21037; Invitrogen) for 1 hr at RT, washed with PBST, and mounted in SlowFade™ Gold Antifade Mountant with DAPI (cat# S36938; Invitrogen). Images were taken using a BZ-X800 All-in-one Fluorescence Microscope (Keyence).

### Statistical analysis

All data are presented as mean ± standard deviation (SD). Two-way ANOVA with the Šídák method for multiple comparisons was performed to analyze the effect of sex and genotype on each parameter. *P*-values less than 0.05 were considered statistically significant. Statistical analyses were carried out using Graph Pad Prism 9.0 software.

## Non-standard Abbreviations and Acronyms

NLS: nuclear localization signal
DCM: dilated cardiomyopathy
RRM: RNA recognition motif
RS: arginine/serine-rich
RBM20: RNA binding motif 20
SR: serine arginine
IHC: immunohistochemistry
LV: left ventricular
AW: anterior wall
PW: posterior wall
FS: fractional shortening
EF: ejection fraction
SDS: sodium dodecyl sulfate
VAGE: vertical SDS-agarose gel electrophoresis
RNA-seq: RNA sequencing
rMATS: multivariate analysis of transcript splicing for replicates
SE: skipped exon
A5SS: alternative 5’ splice site
A3SS: alternative 3’ splice site
MXE: mutually exclusive exon
RI: retained intron
FDR: false discovery rate
ICC: immunocytochemistry
DMEM: Dulbecco’s Modified Eagle Medium
SD: standard deviation
BW: body weight
HW: heart weight
LVID;s: LV inner diameter at end systole
LVID;d: LV inner diameter at end diastole
ESV: end systolic volume
EDV: end diastolic volume
SV: stroke volume
CO: cardiac output
DSGs: differentially spliced genes
N2BA-G: giant N2BA titin isoform
snRNP: small nuclear ribonucleoprotein

## Data availability

Raw and processed data from RNA-Seq are available in the Gene Expression Omnibus (GEO) database under the accession number GSE212799. Source data are provided with this paper. The remaining data supporting the findings of this study are available within the Article, Supplementary Information or Source Data file.

## Acknowledgments

The authors would like to thank the University of Wisconsin-Madison Genome Editing and Animal Models (GEAM) Core Facility for their assistance in generating *Rbm20*^*ΔRS*^ mice. We would also like to thank the University of Wisconsin-Madison Biotechnology Center Gene Expression Center & DNA Sequencing Facility for providing library preparation and next generation sequencing services, and the histology support from the Dairy Innovation Hub Histology Resource. This work was supported by the NIH NHLBI HL148733; the American Heart Association Foundation 19TPA3480072; the Wisconsin Alumni Research Foundation AAH4884 and the University of Wisconsin Foundation AAH5964.

## Contributions

Y.Z., Z.R.G., and Y.L. performed animal studies. Y.Z. and Y.W. carried out biochemical studies. P.L. assisted with biochemical studies. N.A. helped with histology. C.U.B. and J.Z. conducted RNA-seq and bioinformatics analyses. T.A.H. carried out echocardiography. H.G. provided *Rbm20*^*ΔRRM*^ mice. Y.Z., Z.R.G., and W.G. wrote the manuscript. C.U.B., J.Z., T.A.H., and H.G. edited the manuscript. W.G. conceived and supervised the study, analyzed data, and secured funding.

