## Supplemental information for "Disruption of the nuclear localization signal in RBM20 is causative in dilated cardiomyopathy"

**SUPPLEMENTAL MATERIAL**

**Disruption of the nuclear localization signal (NLS) in the RS domain is causative in dilated cardiomyopathy caused by RBM20 mutations**

Yanghai Zhang, MS^1,#^, Zachery R. Gregorich, PhD^1,#^, Yujuan Wang, MS^1^, Camila Urbano Braz, PhD^1^, Jibin Zhang, PhD^2^, Yang Liu, PhD^1^, Peiheng Liu^1^, Nanyumuzi Aori^3,4^, Timothy A. Hacker, PhD^5^, Henk Granzier, PhD^6^, Wei Guo, PhD^1,7,^*

^1^Department of Animal and Dairy Sciences, University of Wisconsin-Madison, Madison, WI 53706, USA

^2^Department of Anatomic Pathology, Comprehensive Cancer Center, City of Hope, Duarte, CA 91010, USA

^3^Department of Kinesiology, University of Wisconsin-Madison, Madison, WI 53706, USA

^4^Department of Psychology, University of Wisconsin-Madison, Madison, WI 53706, USA

^5^Division of Cardiovascular Medicine, Department of Medicine, University of Wisconsin School of Medicine and Public Health, Madison, WI 53706, USA

^6^Department of Cellular and Molecular Medicine, University of Arizona, Tucson, AZ 85724, USA

^7^Cardiovascular Research Center, School of Medicine and Public Health, University of Wisconsin-Madison, Madison, WI 53706, USA

^#^Equal contribution

Short title: Loss of RBM20 nuclear localization causes DCM

Table S1. Single-stranded donor and sgRNAs for CRISPR/Cas9 genome editing.

| **Name** | **Sequence (5’→3’)** |
| --- | --- |
| Single-stranded donor | CCATCTGGGTGATGCAGGTTACGAGCTCTGCAGAGTCTAAACCCTGTCTCTTCCCTTCCTCCCAGGTATGGTCCAtatGCGTGGAGAGACGAGGATCGAGAGACTGTCCCCAGGAGGGAGAACGGGGAAGACAAAAGAGACAGGTTGGATGTT |
| sgRNA 1 | ATGGCCGTGACTCCTACGCG |
| sgRNA 2 | CTCCCAGGTATGGTCCAGAG |

Table S2. Primer information.

| **Experiment** | **Gene** | **Sequences (5'→3')** | **Annealing temp.** | **Products (bp)** |
| --- | --- | --- | --- | --- |
| Genotyping | *Rbm20* | F: TTCCTGGACACTCTGCACTAC  R: CTCATCCAACTCAGCTTTGTCC | 63 ^o^C | WT: 405  ΔRS: 303 |
| RT-PCR | *Camk2d* | F: CGAGAAATTTTTCAGCAGCC  R: GTCTTCATCCTCAATGGTGGTG | 60 ^o^C | 197, 155, 128, 95 |
| RT-PCR | *Ryr2* | F: GTTGTCACGATGAAGAAGACGATG  R: CTTTGCTGGCACTGATAGTCTG | 60 ^o^C | 177, 153 |
| RT-PCR | *Tpm2* | F: AGAGCCGAGGTGGCTGA  R: TCAGCTTCTCCTCCAGAAGT | 60 ^o^C | 154, 230 |
| RT-PCR | *Gapdh* | F: GGTGGACCTCATGGCCTACA  R: CTCTCTTGCTCAGTGTCCTTGCT | 60 ^o^C | 82 |

‘F’ and ‘R’ denote forward and reverse primers, respectively. ‘ΔRS’ indicates the band size in gene edited *Rbm20^ΔRS^* mice lacking 102 bp stretch in the *Rbm20* gene corresponding to the RS domain.

Table S3. Cardiac structure and function in 4-month-old male WT (n=8) and *Rbm20^ΔRS^* (n=7) mice, as well as female WT (n=7) and *Rbm20^ΔRS^* (n=11) mice as assessed by M-mode echocardiography.

|  |  | **Male** | | | **Female** | | | **Two-way ANOVA (*p*-value)** | | |
| --- | --- | --- | --- | --- | --- | --- | --- | --- | --- | --- |
| **Parameter** | **Units** | **WT** | **ΔRS** | **Adj. *p*-value** | **WT** | **ΔRS** | **Adj. *p*-value** | **Sex** | **Genotype** | **Interaction** |
| Heart Rate | BPM | 474.75 ± 50.04 | 536.30 ± 81.36 | 0.1787 | 512.72 ± 43.95 | 535.88 ± 71.81 | 0.7407 | 0.4462 | 0.0920 | 0.4360 |
| LVID;s | mm | 3.27 ± 0.31 | 4.08 ± 0.19 | <0.0001 | 2.88 ± 0.29 | 3.85 ± 0.32 | <0.0001 | 0.0098 | <0.0001 | 0.4752 |
| LVID;d | mm | 4.29 ± 0.29 | 4.77 ± 0.11 | 0.0008 | 3.89 ± 0.20 | 4.45 ± 0.21 | <0.0001 | 0.0002 | <0.0001 | 0.6510 |
| ESV | uL | 43.67 ± 10.46 | 73.59 ± 8.32 | <0.0001 | 32.36 ± 8.68 | 64.81 ± 12.28 | <0.0001 | 0.0157 | <0.0001 | 0.7496 |
| EDV | uL | 83.06 ± 13.34 | 105.87 ± 5.79 | 0.0006 | 65.99 ± 8.51 | 90.36 ± 10.14 | 0.0001 | 0.0002 | <0.0001 | 0.8374 |
| Stroke Volume | uL | 39.39 ± 4.55 | 32.28 ± 7.50 | 0.0329 | 33.63± 2.52 | 25.56 ± 4.70 | 0.0088 | 0.0028 | 0.0004 | 0.8037 |
| Ejection Fraction | % | 47.92 ± 4.39 | 30.47 ± 6.85 | <0.0001 | 51.65 ± 6.33 | 28.84 ± 7.48 | <0.0001 | 0.6709 | <0.0001 | 0.2800 |
| Fractional Shortening | % | 23.93 ± 2.50 | 14.44 ± 3.64 | <0.0001 | 26.11 ± 3.72 | 13.50 ± 3.91 | <0.0001 | 0.6446 | <0.0001 | 0.2493 |
| Cardiac Output | mL/min | 18.70 ± 2.91 | 16.99 ± 3.13 | 0.3825 | 17.27 ± 2.15 | 13.44 ± 1.61 | 0.0098 | 0.0114 | 0.0054 | 0.2598 |
| LV Mass | mg | 91.56 ± 14.88 | 107.41 ± 11.11 | 0.0432 | 64.05 ± 5.43 | 76.29 ± 12.76 | 0.1061 | <0.0001 | 0.0039 | 0.6897 |
| LV Mass Cor | mg | 3.27 ± 0.45 | 3.98 ± 0.35 | 0.0429 | 2.96 ± 0.32 | 3.62 ± 0.74 | 0.0435 | 0.1105 | 0.0019 | 0.9046 |
| LVAW;s | mm | 0.75 ± 0.05 | 0.70 ± 0.05 | 0.3032 | 0.72 ± 0.09 | 0.60 ± 0.03 | 0.0011 | 0.0084 | 0.0009 | 0.1189 |
| LVAW;d | mm | 0.58 ± 0.05 | 0.60 ± 0.04 | 0.8457 | 0.54 ± 0.09 | 0.51 ± 0.06 | 0.5782 | 0.0065 | 0.7906 | 0.3127 |
| LVPW;s | mm | 0.77 ± 0.10 | 0.70 ± 0.08 | 0.2589 | 0.71 ± 0.08 | 0.58 ± 0.07 | 0.0097 | 0.0069 | 0.0034 | 0.3395 |
| LVPW;d | mm | 0.62 ± 0.09 | 0.58 ± 0.07 | 0.5075 | 0.50 ± 0.07 | 0.47 ± 0.06 | 0.6518 | 0.0001 | 0.1891 | 0.8403 |

LVID;s, LV inner diameter during systole; LVID;d, LV inner diameter during diastole; ESV, end systolic volume; EDV, end diastolic volume; EF, ejection fraction; FS, fractional shortening; LVAW;s, LV anterior wall thickness at end of systole; LVPW;s, LV posterior wall thickness at end of systole. Data presented as mean ± SD. Two-way ANOVA with the Šídák method for multiple comparisons was performed to analyze the effect of sex and genotype on each individual parameter.

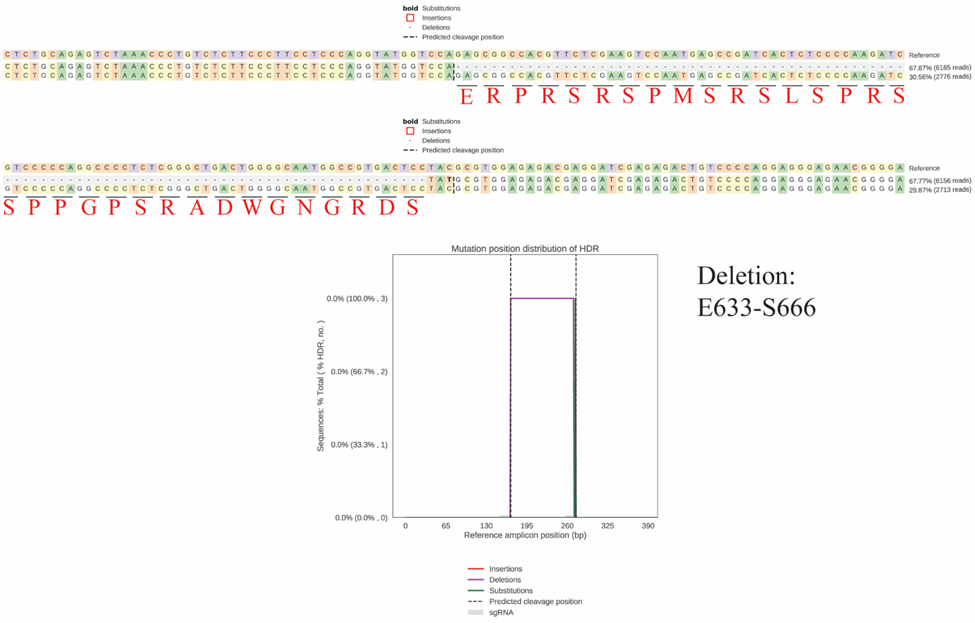

**Figure S1. Confirmation of CRISPR/Cas9 genome edits in heterozygous founder via deep sequencing using an Illumina MiSeq System.** The amino acid sequence of the 102 bp stretch deleted in the gene edited allele is labeled below the corresponding DNA sequence.

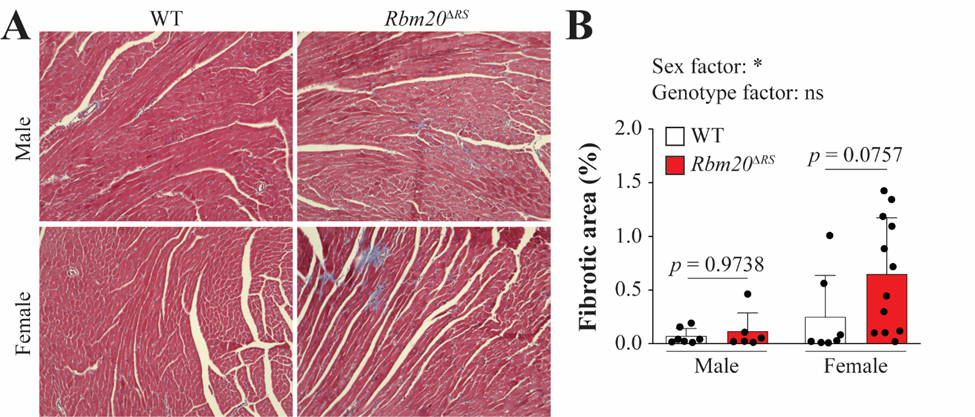

**Figure S2. Quantification of fibrosis in the hearts of 4-month-old male WT (n=7) and *Rbm20^ΔRS^* (n=6) mice, as well as female WT (n=7) and *Rbm20^ΔRS^* (n=12) mice.** **A**, Representative images showing Masson’s trichromed stained tissue sections from male and female WT and *Rbm20^ΔRS^* mice. **B**, Quantification of the fibrotic area in the hearts of WT and *Rbm20^ΔRS^* mice of both sexes. Data presented as mean ± SD. Two-way ANOVA with the Šídák method for multiple comparisons was performed to analyze the effect of sex and genotype on each individual parameter. ns, not significant; **p*<0.05.

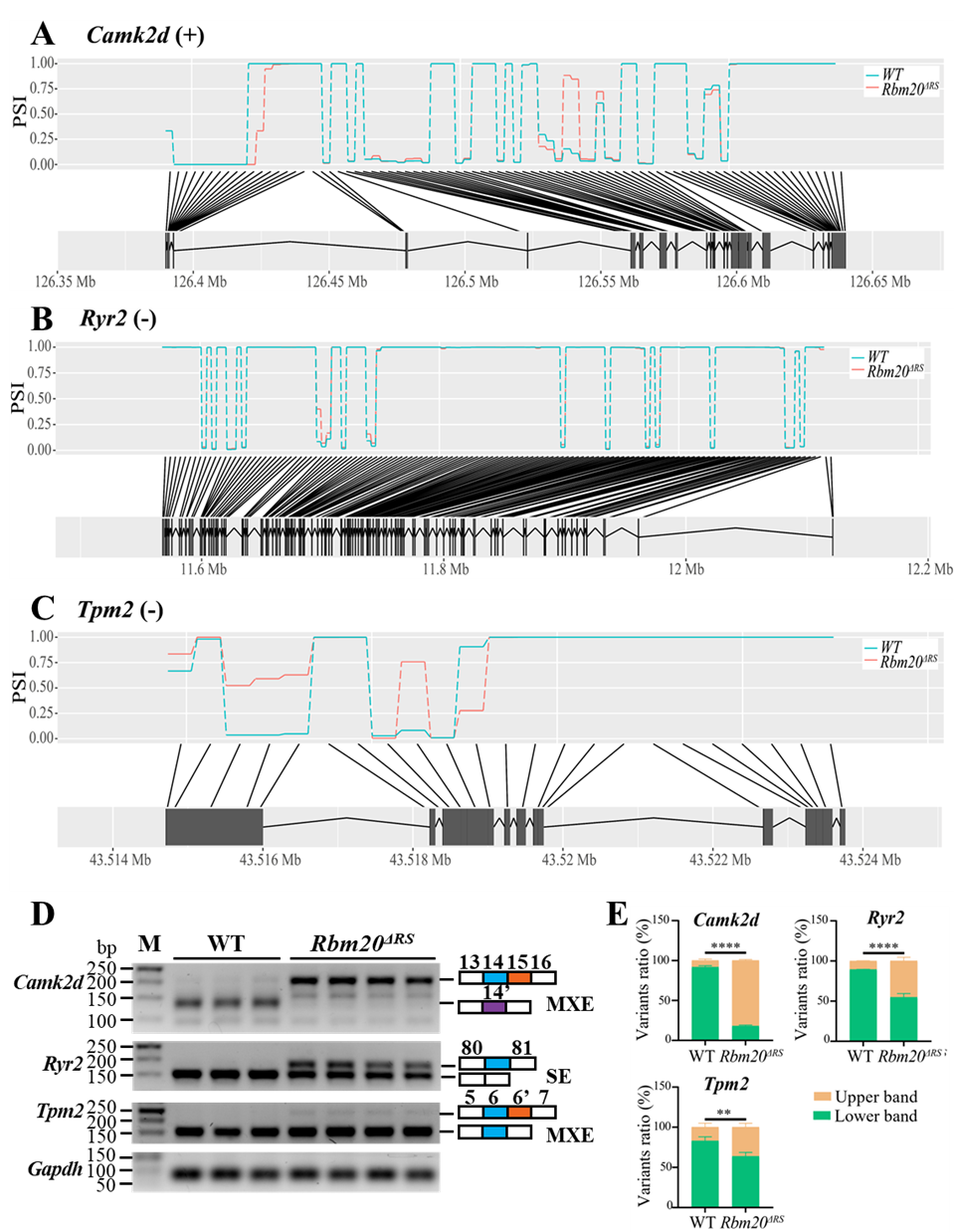

**Figure S3. Validation of RBM20 target transcript splicing changes in the hearts of 2-month-old male *Rbm20^ΔRS^* mice. A-C**, RNA-seq PSI alternative splicing maps for *Camk2d* (**A**), *Ryr2* (**B**), and *Tpm2* (**C**) comparing WT (blue) and *Rbm20^ΔRS^* (red). **D**, RT-PCR validation of *Camk2d*, *Ryr2*, and *Tpm2* alternative splicing in the hearts of *Rbm20^ΔRS^* mice. **E**, Quantitative analysis of RT-PCR results. The variant ratio was calculated as the intensity of the upper band versus the intensity of the lower band for each gene. Data are presented as mean ± SD. ns, not significant; **p*<0.05; ***p*<0.01; ****p*<0.001; *****p*<0.0001.

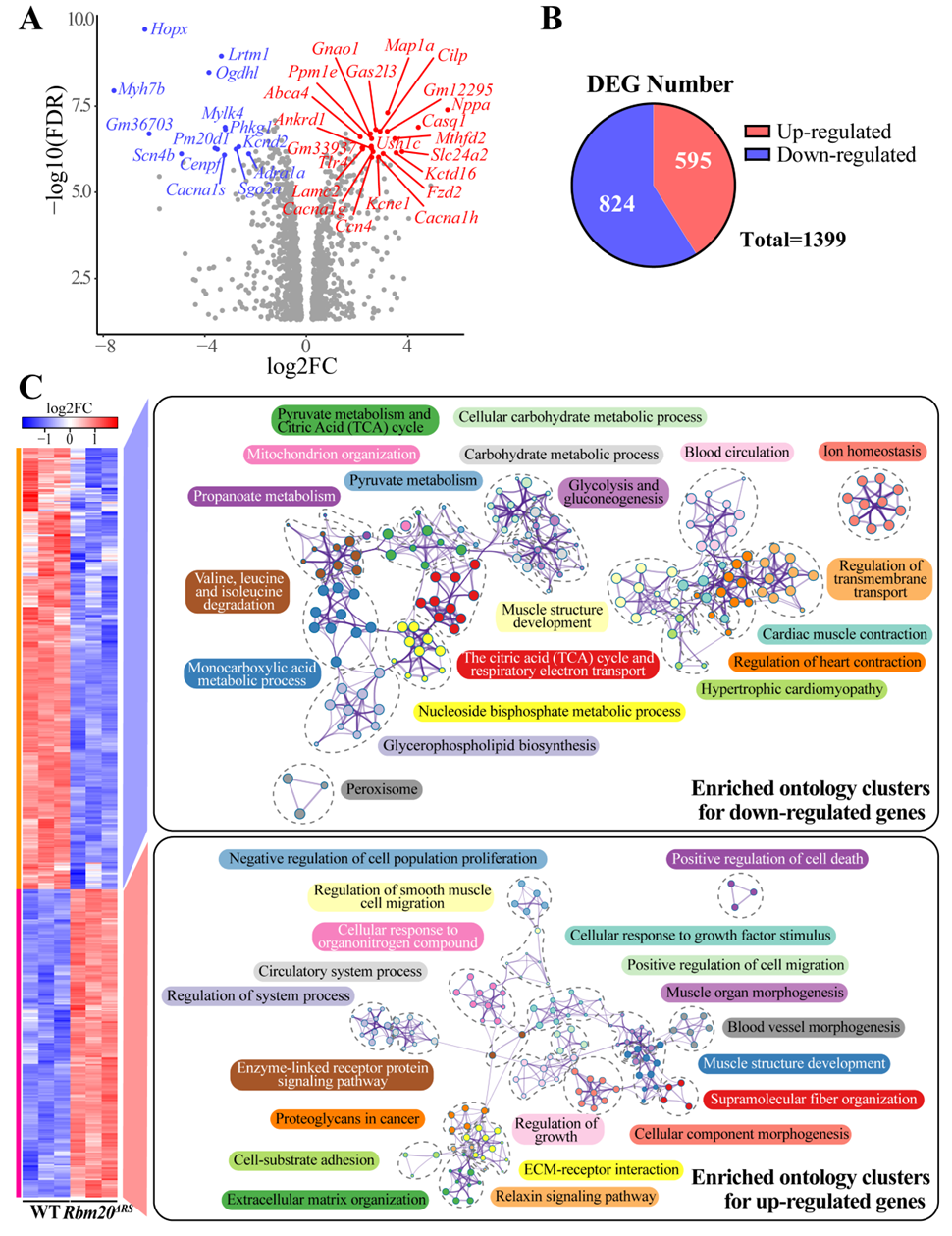

**Figure S4. Analysis of genes differentially expressed in the hearts of 2-month-old male *Rbm20^ΔRS^* mice relative to age- and sex-matched WT controls. A**, Volcano plot showing genes differentially expressed in the hearts of *Rbm20^ΔRS^* relative to WT control mice. Log2 fold change (log2FC) was plotted against the -log10(FDR) value. Genes with -log10(FDR) > 6 and |log2FC| > 2 are indicated in red and blue, respectively. **B**, Pie chart showing the total number of up- and down-regulated genes in *Rbm20^ΔRS^* mouse hearts versus WT. **C**, Heatmaps comparing gene expression in the hearts of *Rbm20^ΔRS^* and WT mice. Genes down- or up-regulated in *Rbm20^ΔRS^* mice versus WT were analyzed using Metascape and the top 20 enriched ontology clusters are shown with their representative enriched terms listed (one per cluster).
